# Genetic disruption of cutaneous de novo glucocorticoid synthesis: a CRISPR-Cas9 CYP11B1-knockout human epidermal model

**DOI:** 10.64898/2026.11.03.764005

**Authors:** Sihui Xu, Liu Yang, Kunal Das Mahapatra, Guanglin Niu, Onur Parlak

## Abstract

Human skin functions as both a physical barrier and a neuroendocrine and steroidogenic organ. Glucocorticoids (GC) synthesized in the skin help to maintain skin immune homeostasis and to mediate responses to environmental stressors. Cutaneous GC synthesis has so far been disrupted only pharmacologically, with inhibitors that do not discriminate CYP11B1 from its close paralogue CYP11B2, or genetically in mouse keratinocytes, so the consequences of blocking this pathway in human keratinocytes remain untested. Here, we disrupted de novo GC synthesis by knocking out the CYP11B1 gene using CRISPR-Cas9, thereby interrupting the terminal enzymatic process of converting 11-deoxycortisol to cortisol in mitochondria. Two independent knockout clones were established, one carrying a 28 bp deletion spanning the start codon and one carrying a multi-exon deletion. Both clones reconstituted stratified 3D epidermal models as in wild-type models, indicating that loss of CYP11B1 does not by itself impair epidermal differentiation. Wild-type models responded with increased CYP11B1 transcript abundance and immunofluorescence signal, whereas both knockout clones retained a UVB-inducible transcript signal but showed no detectable CYP11B1 protein, confirming loss of the protein product at the tissue level. This work provides, to our knowledge, the first human keratinocyte model carrying a genomically defined block in de novo GC synthesis, and a tissue-level platform for resolving the role of keratinocyte-derived GC in epidermal homeostasis and environmental stress responses.

## 1. Introduction

Skin, as one of the largest organs of the human body, is not only a physical barrier but also functions as a neuroendocrine and steroidogenic organ. The epidermis consists mainly of keratinocytes and forms a physical barrier that protects against mechanical forces, limits trans-epidermal water loss and helps to maintain body temperature ^[1,2]^. As an endocrine organ, it also recognizes the environmental stressors through different receptors and generates local neuroendocrine signals, including hormones, molecules, and neuropeptides ^[3–6]^. Recent studies showed that keratinocytes synthesize de novo glucocorticoids (GC) under the hormonal control of the local skin-specific hypothalamic–pituitary–adrenal (HPA) axis ^[7–11]^. In particular, skin-derived GCs regulate skin homeostasis and inflammation, decrease keratinocyte proliferation, promote epidermal differentiation and impair wound healing ^[4,12–16]^.

The de novo GC synthesis is initiated by cholesterol transport into mitochondria, followed by a series of steroidogenic reactions involving various metabolic intermediates ^[3,17–19]^. As a member of the cytochrome P450 family of enzymes, the 11β-hydroxylase (CYP11B1) catalyzes the terminal enzymatic step of synthesizing the bioactive GC that hydroxylates 11-deoxycortisol into cortisol. Downstream of synthesis, cortisol is regenerated from cortisone by 11β-hydroxysteroid dehydrogenase 1 (11β-HSD1, encoded by HSD11B1) in the endoplasmic reticulum and is inactivated to cortisone by 11β-hydroxysteroid dehydrogenase 2 (11β-HSD2, encoded by HSD11B2). Therefore, as a key enzyme, the expression of CYP11B1 in keratinocytes illustrates the capability of de novo GC synthesis in the skin ^[8,14]^. Furthermore, environmental stressors such as ultraviolet radiation (UV) have been shown to be associated with the activation of the skin-specific HPA axis and to enhance de novo GC synthesis ^[20–23]^. As described by Skobowiat *et al*., the stimulation level of CYP11B1 is dependent on UV wavelengths, which are UVA (320–400 nm), UVB (280–320 nm), and UVC (100–280 nm) ^[24]^. Among these, UVB was found to upregulate CYP11B1 expression in human skin, with the magnitude of induction increasing with UVB dose.

Although increasing evidence supports GC synthesis in the epidermis, the specific consequences of disrupting de novo GC synthesis in human keratinocytes remain unclear. Previous studies relied on the CYP11B1 inhibitor metyrapone to interrupt cortisol synthesis in the skin ^[4,25,26]^. Pharmacological inhibition is, however, reversible and dose-dependent, and metyrapone additionally inhibits 11β-HSD1, so the cellular consequences of inhibitor treatment cannot be assigned unambiguously to the loss of CYP11B1 alone. A second, genetic approach used an *in-vivo* animal model, in which *Cyp11b1* was specifically deleted in mouse keratinocytes to study how skin-derived GC regulates skin immune homeostasis ^[14]^. That work established the immunological importance of keratinocyte-derived GC, but murine and human epidermis differ in differentiation kinetics, barrier composition and steroidogenic enzyme regulation, so the findings cannot be transferred directly to human skin. Although several studies have provided important insights into cutaneous GC biology ^[27,28]^, to our knowledge no human keratinocyte line carrying a heritable, gene-level block in de novo GC synthesis has been reported, and there is correspondingly little evidence on how endogenous GC production in human keratinocytes affects downstream biological processes such as proliferation, differentiation, barrier function, and immune responses. Therefore, establishing an epidermal model with a well-defined genetic background and stable characteristics, in which the de novo GC synthesis pathway is disrupted, would provide a valuable experimental platform for better understanding the biological functions of skin-derived GC.

This can be achieved by using CRISPR-Cas9 genome editing to knock out (KO) the CYP11B1 gene, thereby disrupting de novo GC synthesis in human keratinocytes. The CRISPR-Cas9 system is widely used to introduce defined gene mutations by inducing targeted DNA double-strand breaks, which are subsequently repaired through non-homologous end joining (NHEJ)-mediated repair or homology-directed repair (HDR) pathways ^[29–31]^. The NHEJ-mediated DNA repair induces small insertion or deletion mutations (indels), which cause frameshifts and can introduce premature stop codons ^[32,33]^. In addition, targeting more than one genomic site across several exons can generate a large deletion, which can increase KO efficiency for the target gene ^[34,35]^. Moreover, compared to primary keratinocytes, the immortalized cell lines are more widely used for CRISPR-Cas9 genome editing because they are morphologically stable over passages, have an extended or unlimited lifespan, and allow edited monoclonal lines to be established more readily ^[36,37]^. Among them, the immortalized human N/TERT-2G keratinocyte cell line has been shown to be well suited to the generation of CRISPR-Cas9 KO models. An additional design consideration for this locus is that CYP11B1 shares approximately 93% coding-sequence identity with its paralogue CYP11B2 (aldosterone synthase), so it is difficult to make gene-specific mutations at this locus, and guide RNAs must be selected in regions that discriminate between the two genes ^[38–40]^.

Although common two-dimensional (2D) keratinocyte cultures are valuable for evaluating genome editing and molecular genotype changes, these methods fail to replicate key characteristics of human epidermis such as proliferation, differentiation, and stratum corneum formation ^[41–43]^. Three-dimensional (3D) reconstructed epidermal models provide a more physiologically relevant *in-vitro* environment by reproducing key features of natural epidermal structure and biological function ^[44–47]^. Although several immortalized keratinocyte cell lines differentiate poorly, the N/TERT-2G cell line shows morphology and differentiation architecture similar to those of human primary keratinocytes ^[48,49]^. Therefore, developing 3D epidermal models using the CYP11B1-KO N/TERT-2G keratinocytes enables evaluation of the impact of de novo GC synthesis deficiency on epidermal differentiation, barrier function, and homeostasis at the tissue level.

In this study, we set out to close that gap by establishing what is, to our knowledge, the first genetically defined human epidermal model in which keratinocyte-derived de novo GC synthesis is disrupted at the genomic level rather than pharmacologically. To do so, we used CRISPR-Cas9 to knockout the CYP11B1 gene in N/TERT-2G keratinocytes, followed by genomic and molecular characterization. We then used these cells to fabricate 3D epidermal models to evaluate how disrupted de novo GC synthesis affects epidermal differentiation and phenotype. UVB exposure was used as an environmental stressor to stimulate the expression of CYP11B1, thereby validating the gene-editing result of CYP11B1 knockout. Because the disruption is heritable and gene-specific, the resulting CYP11B1-KO 3D model complements inhibitor-based and murine approaches with a stable human system, and provides a genetically defined platform for understanding the role of de novo synthesized GC in epidermal homeostasis and environmental stress responses.

## 2. Results

### 2.1. CYP11B1 KO keratinocytes generation

To generate the CYP11B1 KO N/TERT-2G keratinocytes, we designed two CRISPR-Cas9 editing strategies: indels targeting the start codon and a large deletion between exons.

#### Strategy I: Indels targeting the start codon

Two CRISPR RNAs (crRNAs) were designed to target the start codon (ATG) region of the CYP11B1 gene (Figure 1A, B; Table S1) and annealed with tracrRNA to form two-part guide RNAs (gRNAs) to improve editing efficiency. Then, the gRNAs were assembled with Cas9 into ribonucleoprotein (RNP) complexes and introduced into N/TERT-2G keratinocytes via electroporation. The RNP complexes induced double-strand DNA breaks, which the cells repaired through the NHEJ mechanism, resulting in indel mutations. Following cell culture, genomic DNA was extracted from the gene-edited cell pool and amplified using designed PCR primers; the expected amplification product size for the wild-type (WT) DNA sequence was 401 bp (Figure S1A; Table S2). The amplified DNA fragments were subjected to Sanger sequencing and subsequently analyzed against the WT sequence using the Inference of CRISPR Edits (ICE) tool ^[50]^ to determine the indel editing outcomes and efficiency (Figure S1B, C). The cell pool exhibiting the highest editing efficiency was then selected for single-cell culture and serial expansion to establish monoclonal cell populations. To validate the CRISPR-Cas9 genome editing and clonality of the CYP11B1 KO cell lines, we extracted genomic DNA from each monoclonal population and amplified it using the same PCR primers (Table S2). Subsequently, PCR-amplified DNA fragments were sequenced through Sanger sequencing, and the results compared between the CYP11B1 KO monoclonal populations (data not shown) and WT keratinocytes using Benchling. Sequencing revealed a 28 bp deletion spanning the CYP11B1 start codon (ATG) in the B5 clone (Figure 1C, S2A). In addition, the ICE tool was used to analyze indels and their relative prevalence for the B5 clone; the analysis returned an indel frequency of 100%, indicating that no wild-type allele was retained in this clone (Figure 1D). Consistent with this, agarose gel electrophoresis showed that the amplicon from the B5 clone was smaller than that from WT keratinocytes (Figure 1E).

**Figure 1.**
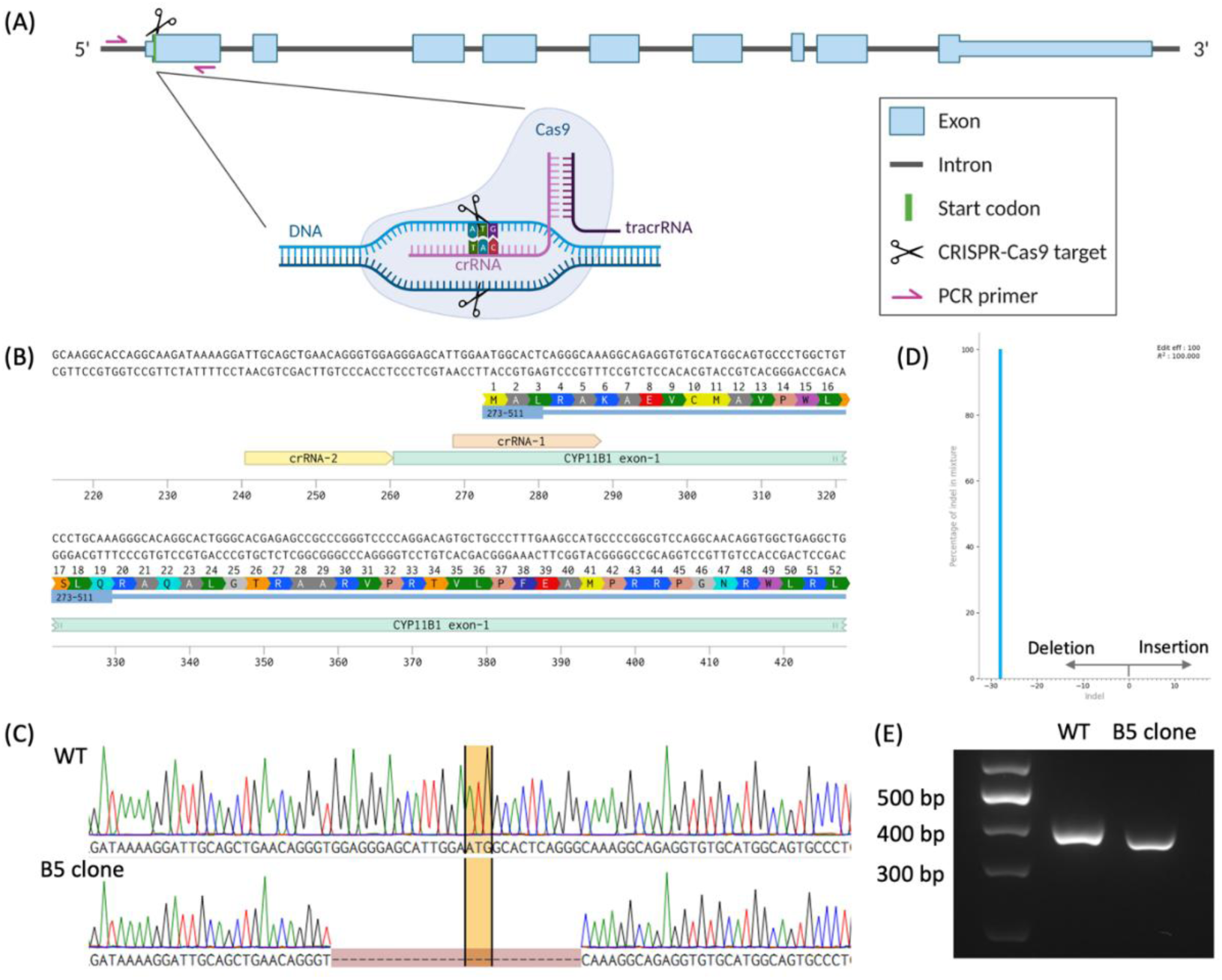
*Strategy I:* indels targeting the start codon (ATG) of CYP11B1. (A) Schematic of the CRISPR-Cas9 complex target and PCR primer locations in CYP11B1 sequence. (B) Locations of the CRISPR RNAs (crRNAs) designed to disrupt the start codon (ATG) of CYP11B1 in N/TERT-2G keratinocytes. (C) PCR amplified sequence of B5 clone, aligned with WT through Benchling, indicates a 28 bp deletion at the ATG site of CYP11B1. (D) Indels and their relative prevalence for the B5 clone, calculated by ICE. (E) PCR amplification and agarose gel electrophoresis of the CYP11B1 target region from genomic DNA of WT keratinocytes and the B5 clone. WT: wild-type N/TERT-2G keratinocytes, B5: CYP11B1 KO monoclonal cell line.

#### Strategy II: A large deletion between exons

Three pre-designed Alt-R CRISPR-Cas9 crRNAs targeting different exons of CYP11B1 were obtained from Integrated DNA Technologies (IDT) to induce a large-fragment deletion (Figure 2A; Table S1). The CRISPR-Cas9 genome editing workflow followed *strategy I*, involving RNP complexes preparation and electroporation delivery into N/TERT-2G keratinocytes. After 48 hours of culture, the cells were detached and subjected to single-cell culture and expansion. The first round of screening was then performed: genomic DNA was extracted from each monoclonal cell population and amplified using designed PCR primers targeting the intron region between exon 2 and exon 3; the amplification product size of WT was 836 bp (Figure 2A; Table S2). Comparison of the PCR-amplified DNA fragments via agarose gel electrophoresis revealed distinct DNA bands at 836 bp in both N/TERT-2G WT keratinocytes and primary keratinocytes (Figure 2B). Based on the absence of this DNA band in the agarose gel electrophoresis results, three CYP11B1 KO candidate monoclonal clones (SA2, SC1, and DB4) were selected. Subsequently, a second round of screening was conducted using designed PCR primers flanking the CRISPR-Cas9 mediated cleavage region (Figure 2C; Table S2). For the CYP11B1 KO monoclonal cell populations, the PCR product size was approximately 600 bp; in contrast, the 3914 bp product size expected for WT keratinocytes was not amplified under the conditions used (Figure 2D). The amplified DNA fragments were sent for Sanger sequencing, and the results were analyzed by aligning with the WT DNA sequence using Benchling. Ultimately, the DB4 clone was selected on the basis of the larger deletion (approximately 3300 bp, removing exons 1-5; Figure S2B), together with its cellular morphology and population doubling time.

**Figure 2.**
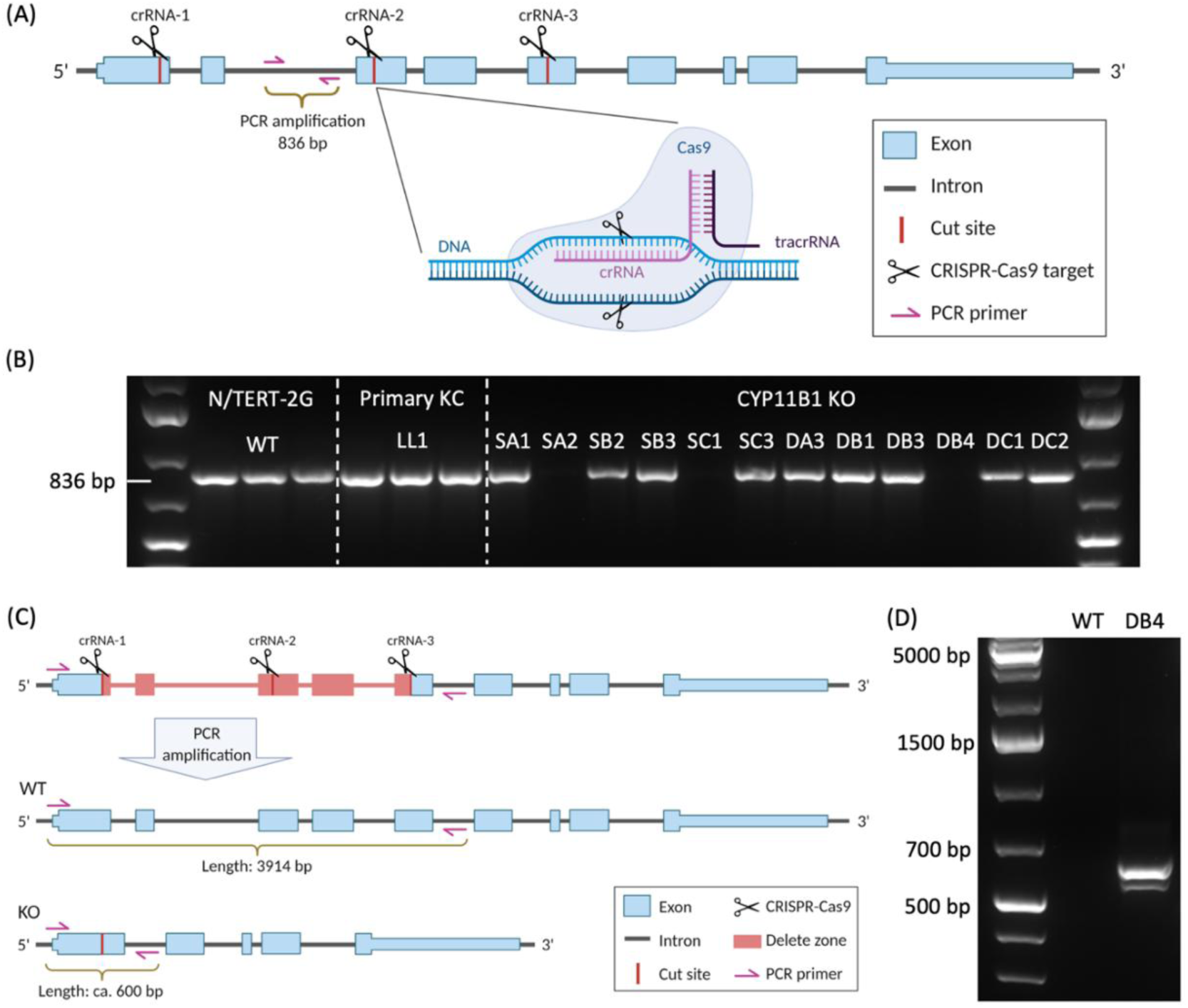
*Strategy II:* a large deletion between CYP11B1 exons. (A) Schematic of the CRISPR-Cas9 complex target locations on CYP11B1 DNA sequence, and PCR primers for the first round of screening. (B) PCR amplification and agarose gel electrophoresis of the extracted genomic DNA of WT keratinocytes, primary keratinocytes (LL1), and CYP11B1 KO monoclonal populations. (C) Schematic of the PCR primers for the second round of screening, and the estimated DNA fragments of WT keratinocyte and CYP11B1 KO monoclonal population. (D) PCR amplification and agarose gel electrophoresis of the extracted genomic DNA of WT and DB4 clone. WT: wild-type N/TERT-2G keratinocytes, DB4: CYP11B1 KO monoclonal cell line.

### 2.2. Establishment and validation of 3D epidermal model with KO keratinocytes

To further understand the effect of CYP11B1 KO on epidermal differentiation, barrier function, and homeostasis at the tissue level, we utilized the CYP11B1 KO monoclonal cell lines and WT N/TERT-2G keratinocytes to fabricate 3D epidermal models. As described in our previous study ^[47]^, keratinocyte suspensions were seeded separately onto collagen pre-coated cell culture inserts in 6-well plates and cultured for three days, allowing the keratinocytes to proliferate and reach confluence on the insert membranes. Subsequently, the medium inside the insert was removed to generate an air–liquid interface over the keratinocyte monolayer and initiate differentiation. After 12 days of culture, the 3D epidermal models were harvested for further study.

To validate the expression of several key epidermal differentiation biomarkers in 3D epidermal models generated from CYP11B1 KO monoclonal cell lines (B5 and DB4 clones) and WT keratinocytes, the paraffin-embedded tissue sections were immunofluorescent stained for keratin 14 (K14, basal layer keratinocyte proliferation), keratin 10 (K10, suprabasal layer keratinocyte differentiation), filaggrin (FLG, terminal differentiation), and claudin-1 (CLDN1, tight junction). Immunofluorescence results demonstrated that the differentiation characteristics of 3D epidermal models derived from immortalized N/TERT-2G keratinocytes were largely consistent with those of 3D epidermal models derived from primary human keratinocytes and *ex-vivo* human skin samples ^[47]^, as evidenced by the clear presence of all these biomarkers in the corresponding locations of the epidermis (Figure 3A, S3). Furthermore, the B5 and DB4 clones exhibited similar differentiation profiles, indicating that deletion of CYP11B1 did not produce apparent differences in the distribution of these differentiation markers under the conditions tested.

**Figure 3.**
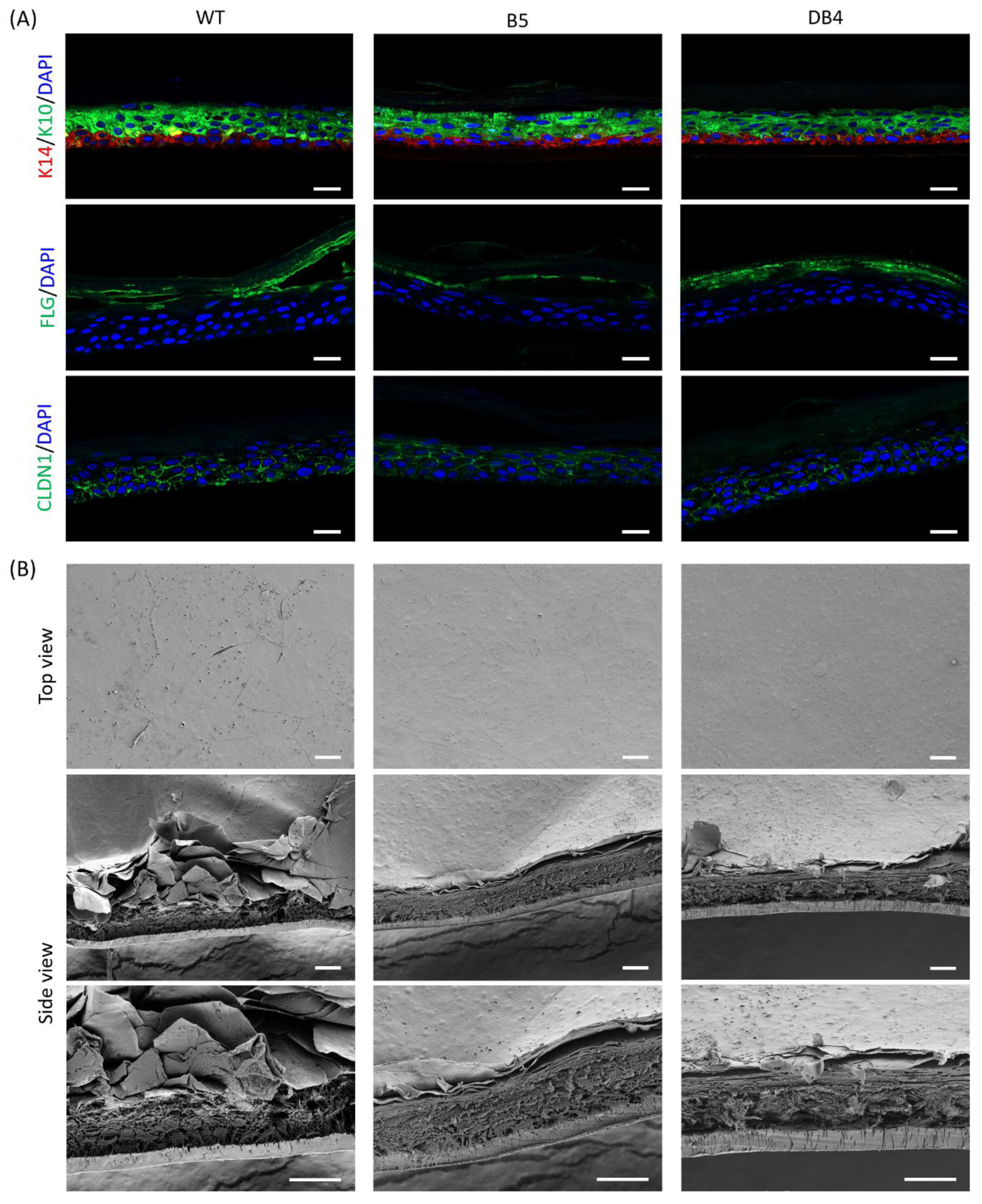
Differentiation and morphological validation of the 3D epidermal models. (A) Immunofluorescent stains on the paraffin-embedded 3D epidermal sections of key biomarkers representing keratinocyte differentiation in human epidermis. (B) Scanning electron microscopy images of the 3D epidermal models. WT: wild-type N/TERT-2G keratinocytes; B5 and DB4: CYP11B1 KO monoclonal lines. Scale bars: 20 μm.

Subsequently, we verified the morphological structure of 3D epidermal models using scanning electron microscopy (SEM). Compared to 3D epidermal models derived from primary keratinocytes, those generated from immortalized N/TERT-2G keratinocytes displayed a flatter stratum corneum in top-view images, although a distinct scaly cornified layer remained visible in side-view images (Figure 3B). Similar trends were also observed in both B5 and DB4 clones.

### 2.3. CYP11B1 KO validation through UVB irradiation

As described above, CYP11B1 expression is induced by stressors such as ultraviolet radiation, enabling the terminal mitochondrial step of cortisol synthesis. In this study, UVB was used as an environmental stressor to induce CYP11B1 expression in 3D epidermal models, aiming to validate the success of CYP11B1 KO in N/TERT-2G keratinocytes. UVB irradiation was performed after 12 days of culture, ensuring adequate differentiation of the 3D epidermal models, and samples were collected 24 hours post-treatment for subsequent analysis. We used immunofluorescence staining to detect the CYP11B1 expression in paraffin-embedded tissue sections, comparing control 3D epidermal models (Ctl; not exposed to UVB) with UVB-treated models (exposed to a UVB dose of 1000 mJ/cm²) (Figures 4A and S4). Sections from the WT control group exhibited a faint CYP11B1 expression signal, whereas the sections from the WT UVB-treated group displayed a brighter, more distinct fluorescent signal. Meanwhile, quantitative analysis of immunofluorescence intensity showed a statistically significant increase in fluorescence in WT UVB-treated 3D epidermal models compared with WT control 3D epidermal models (Figure 4C). These results indicate that UVB irradiation induces CYP11B1 protein accumulation in N/TERT-2G 3D epidermal models. In contrast, no obvious fluorescent signal was observed in either B5 clone or DB4 clone sections from the control group or after the UVB treatment, where the fluorescence intensity was comparable to that of the negative staining results (Figure S5).

**Figure 4.**
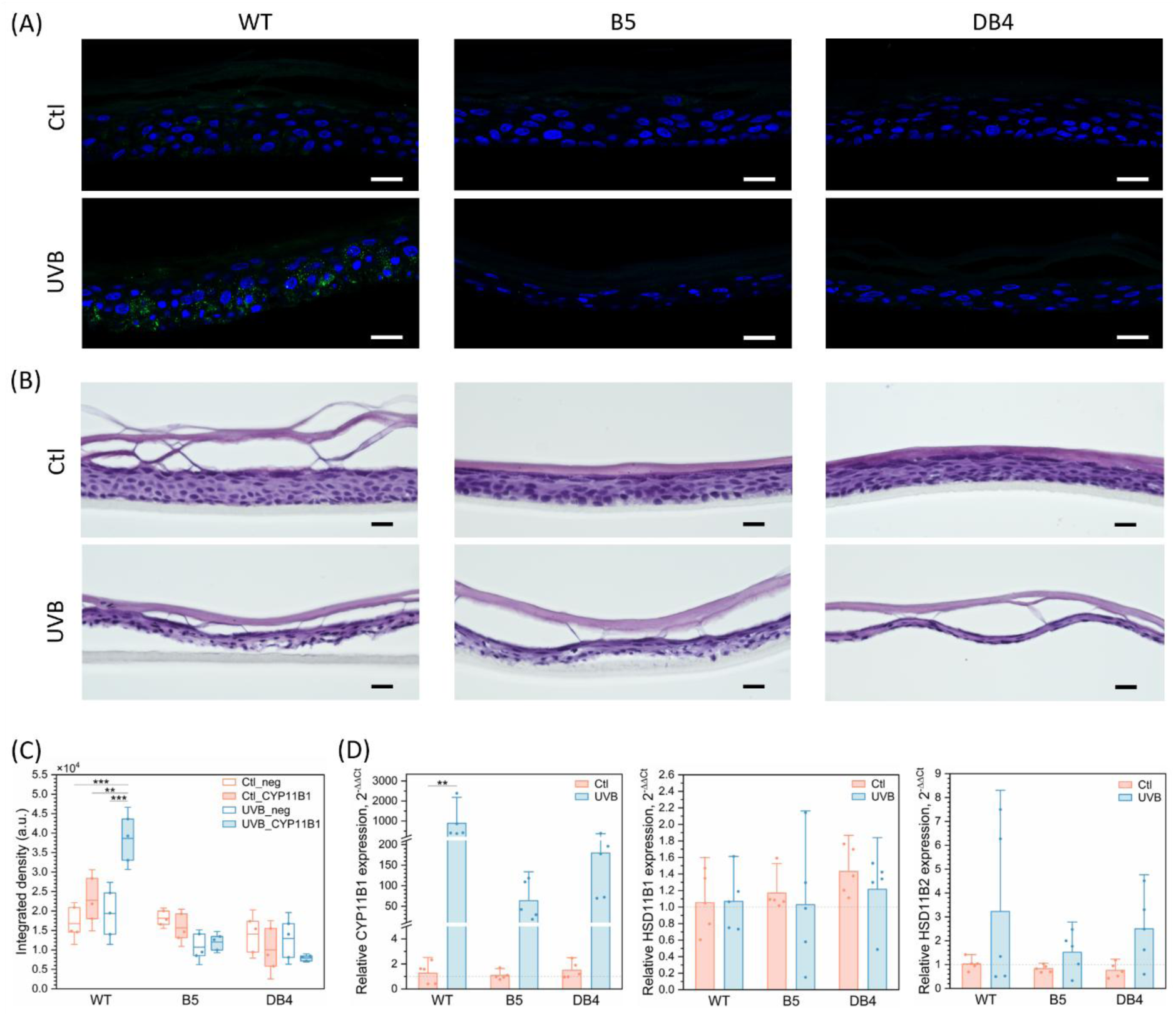
UVB irradiation effect on 3D epidermal models, compared with WT and CYP11B1 KO monoclonal (B5 and DB4 clones) groups. (A) CYP11B1 immunofluorescent staining on the paraffin-embedded 3D epidermal sections. Blue: DAPI, Green: CYP11B1. (B) Histological images of the paraffin-embedded 3D epidermal models. Ctl: control group without any treatment; UVB: exposed to a UVB dose of 1000 mJ/cm². Scale bars: 20 μm. (C) Integrated density of CYP11B1 immunofluorescence (n = 3 independently generated models per group; mean ± SD). (D) Transcription of cortisol synthesis and activation-related genes. Relative mRNA levels were measured by RT-qPCR, normalized to GAPDH and expressed as 2^−ΔΔCt^ relative to the WT control (n = 5 independently generated models per group; mean ± SD). Statistical significance was assessed by paired comparison through *Tukey’s* test; \*\**P* ≤ 0.01, \*\*\**P* ≤ 0.001.

Morphological changes in the 3D epidermal models after UVB treatment were observed in histological and SEM analyses. Histological images were obtained from hematoxylin and eosin (H&E) staining of paraffin-embedded tissue sections to observe the morphological differences between 3D epidermal models derived from WT and CYP11B1 KO monoclonal N/TERT-2G cell lines and the morphological changes after UVB treatment (Figure 4B). The control groups of both WT and CYP11B1 KO (B5 and DB4 clones) 3D epidermal models showed comparable basic morphological organization of proliferative and differentiated compartments in the absence of UVB treatment. This suggests that KO of CYP11B1 did not detectably affect epidermal morphology, which was also validated by the same results in the SEM images (Figure 3B). After UVB treatment, all models showed reduced epidermal thickness and smaller nuclei compared with the corresponding controls. Moreover, the epidermal layer was observed to detach from the cell culture insert membranes, while the stratum corneum layer remained intact in the 3D epidermal models. More details were revealed in the SEM images of the 3D epidermal models, which showed the structural morphological damage caused by the UVB treatment beneath the cornified layer (Figure S6).

Transcript levels of the cortisol synthesis gene CYP11B1 and of the cortisol interconversion genes HSD11B1 and HSD11B2 were then measured using reverse transcription quantitative polymerase chain reaction (RT-qPCR) analysis (Figure 4D). Statistically, higher CYP11B1 transcription levels were observed in UVB-treated WT 3D epidermal models compared to WT untreated ones. This increase is consistent with the higher CYP11B1 immunofluorescence intensity measured in sections of the same models (Figure 4A). However, even though the CYP11B1 transcription was also stimulated by UVB treatment in the 3D epidermal models derived from B5 and DB4 clones, CYP11B1 immunofluorescent signals were not observed in the corresponding sections. This discrepancy between transcription and protein expression can be explained by the experimental design: the qPCR amplicon used here is located in exon 9 (Table S4), outside both the start-codon indel mutation of the B5 clone and the deleted region of the DB4 clone, so residual transcripts responding to UVB treatment can still be detected even when no translatable full-length transcript is produced. The absence of immunofluorescence signals in the same models therefore indicate that the editing events disrupt production of the CYP11B1 protein rather than its transcription. Additionally, HSD11B1 expression did not show a clear difference between the WT and CYP11B1 KO groups or between the UVB-treated and untreated groups. For HSD11B2, slightly higher transcript levels were observed in the UVB-treated groups; this may reflect a homeostatic adjustment in the 3D epidermal model and requires further investigation.

## 3. Discussion

Cutaneous de novo glucocorticoid synthesis has until now been accessible experimentally only through pharmacological inhibition or murine gene deletion. In this study, we established two distinct strategies to generate CRISPR-Cas9-edited N/TERT-2G keratinocyte monoclonal cell lines targeting the CYP11B1 gene, giving a human cellular system in which the pathway is interrupted at its terminal step by a defined genomic lesion. We confirmed that these clones are capable of fabricating 3D epidermal models, which largely retain the expression patterns of differentiation biomarkers characteristic of the epidermal layer. In the wild-type (WT) 3D epidermal models, UVB irradiation led to an increase in both CYP11B1 transcript abundance and its immunofluorescence intensity. In contrast, while UVB-responsive transcriptional signals of the CYP11B1 gene remained detectable in the CYP11B1 KO 3D epidermal models, the CYP11B1 immunofluorescence signal was not observed.

The two editing strategies disrupted CYP11B1 at different genomic sites. The B5 clone carried a 28 bp deletion encompassing the translation start codon (ATG), which is expected to prevent normal initiation of translation at the canonical start site. The DB4 clone contained a larger deletion spanning multiple exons (from exon 1 to exon 5), which was expected to compromise the integrity of the coding region. An indel mutation frequency of 100% in the B5 clone (determined by ICE analysis) indicates that no wild-type allele remained, providing genomic support for a CYP11B1 KO. For the DB4 clone, initial screening was performed by verifying the deletion of intronic segments to identify monoclonal cell lines lacking wild-type alleles. Subsequently, amplification of the entire CRISPR-Cas9-edited region enabled allele-specific sequencing, facilitating the selection of monoclonal cell lines characterized by large-scale genomic deletions.

The ability of both the B5 and DB4 monoclonal lines to form a stratified epidermis demonstrates that the CRISPR-Cas9 editing and monoclonal selection and expansion processes did not impede epidermal reconstruction. The observed biomarker distribution of K14, K10, filaggrin, and claudin-1 indicated that key features, including basal layer identity, suprabasal differentiation, terminal differentiation, and the proper localization of junction-associated proteins, were preserved in the 3D epidermal models. These findings are consistent with previous characterizations of N/TERT-2G keratinocytes as a source for reconstructed epidermal models ^[49]^. Compared to the 3D epidermal models derived from primary keratinocytes, those derived from N/TERT-2G keratinocytes exhibited a flatter stratum corneum surface; this further indicates that, while some epidermal characteristics are retained, the structure is not identical to that of native human epidermis. Such variables must be accounted for when investigating the link between impaired CYP11B1 function and tissue phenotype. The absence of a structural phenotype is itself informative for the use of these lines: because editing, monoclonal selection and expansion leave stratification and marker distribution intact, phenotypes observed in subsequent experiments on these models can be attributed to the loss of de novo GC synthesis rather than to generic tissue damage or to clonal artefacts introduced during editing. Furthermore, given the current cell culture conditions, permeability or electrical barrier function assays are still required to assess more subtle effects on epidermal barrier function.

In gene-edited models, the persistence of RT-qPCR signal for CYP11B1 is a notable observation; however, its interpretation must be distinguished from actual protein expression. RT-qPCR can detect residual transcript fragments, which does not necessarily imply the presence of complete, translationally competent coding sequences. Depending on the primer design, these signals originate from transcript regions retained downstream of the edited intervals; the amplicon used here targets exon 9 and therefore lies outside both the B5 start-codon deletion and the DB4 deleted interval. The responsiveness of these signals to UVB indicates that the CYP11B1 promoter remains inducible in the edited clones. Nevertheless, this does not confirm the production of a functional CYP11B1 protein. Furthermore, no immunofluorescence signal for the CYP11B1 protein could be detected in either clone following UVB exposure, indicating that—under the experimental conditions tested—the levels of the protein (as recognized by the antibody) fall below the detection threshold. This finding provides corroborating evidence for the efficacy of the gene knockout.

UVB irradiation served as an effective experimental approach for assessing the inducible expression of CYP11B1. In the WT 3D epidermal models, UVB exposure led to an increase in CYP11B1 transcript levels and enhanced immunofluorescence signals, consistent with the UV-responsive cutaneous steroidogenic pathway previously observed in human skin and in keratinocyte-melanocyte co-cultures ^[24]^. Under identical irradiation conditions, the B5 and DB4 3D epidermal models did not exhibit comparable levels of CYP11B1 immunofluorescent signals, further confirming that, beyond the basal state, protein expression in these models is indeed disrupted. Additionally, UVB irradiation caused structural damage across all three genotypic models, characterized by epidermal thinning, nuclear shrinkage, detachment of the tissue from the culture insert membrane, and observable damage beneath the remaining stratum corneum. These phenomena indicate that UVB irradiation not only induces the expression of CYP11B1 but also triggers a significant tissue response. Given that damage was observed in the WT, B5, and DB4 3D epidermal models, with no apparent quantitative differences in the extent of injury across genotypes, these morphological alterations cannot currently be attributed solely to impaired CYP11B1 function. Furthermore, the persistence of the stratum corneum does not imply that barrier function remains intact following UVB irradiation. Conducting UVB dose-response and time-course analyses will help to distinguish steroidogenic signaling from general tissue damage and will facilitate the identification of experimental conditions suitable for evaluating genotype-specific responses.

The findings regarding HSD11B1 and HSD11B2 further highlighted the distinction between cortisol biosynthesis and local cortisol metabolism. Previous research has established that human keratinocytes possess the capacity to synthesize cortisol and regulate the interconversion between cortisol and cortisone ^[19]^. In the current 3D epidermal models, no significant differences in HSD11B1 transcript abundance were observed across experimental conditions, suggesting the absence of a clear compensatory mechanism at the transcriptional level at the time point sampled. This does not, however, exclude changes in enzyme abundance or activity. Similarly, the slight increase in HSD11B2 transcripts following UVB irradiation warrants further investigation; yet, in the absence of more robust statistical and functional evidence, it could not be concluded that a homeostatic regulatory mechanism is at play. Even if CYP11B1 activity was lost, cells can still obtain cortisol through the conversion of cortisone or by uptake from the culture environment. Consequently, disruption of CYP11B1 is not synonymous with a complete absence of cortisol or a total blockade of the related signaling pathways.

Although the genomic characterization and the absence of immunofluorescence signals in CYP11B1 KO tissue sections support successful gene disruption, its impact on de novo cortisol synthesis remains to be determined through subsequent monitoring of steroid levels. Furthermore, the well-preserved epidermal structure and the maintained expression patterns of differentiation markers indicate that these gene-edited cells are suitable for constructing three-dimensional epidermal models. However, whether the epidermal barrier function has been altered remains to be evaluated. These considerations define the direction for future functional characterization and establish a foundation for using these 3D models to investigate mechanisms of CYP11B1-dependent steroid synthesis in the human epidermis.

In summary, this study established what is, to our knowledge, the first *in-vitro* human epidermal platform in which CYP11B1 is disrupted at the genomic level, providing a stable and gene-specific alternative to inhibitor treatment and to murine *Cyp11b1* deletion for investigating CYP11B1-dependent steroidogenesis in human tissue. Future work incorporating direct steroid quantification and validation of enzymatic activity will be required to determine the extent to which de novo cortisol synthesis is impaired and how such impairment affects epidermal function. While the CYP11B1 KO monoclonal 3D epidermal model employed here does not allow for conclusions regarding impacts on immune function, the model, together with the wild-type and knockout keratinocyte lines from which it is built, provides a defined and transferable resource for exploring steroid synthesis in human epidermis and its role in the response to environmental stress.

## 4. Materials and Methods

### Materials

EpiLife Medium (MEPI500CA), Human Keratinocyte Growth Supplement (HKGS, S0015), Trypsin-EDTA (0.25%; 25200056), DMEM (41965039), Fetal Bovine Serum (FBS; A5670801), 1× DPBS (14190169), TRIzol Reagent (15596018), Chloroform (158210025), GlycoBlue (AM9516), Agarose (16500100), and 50× TAE Buffer (BP13321) were obtained from Thermo Fisher Scientific. Millicell® cell culture inserts (PIHP01250), CaCl₂ solution (21115), type-1 rat tail collagen (C3867), Vitamin C (L-Ascorbic acid, A5960), and 2-Propanol (1096342500) were purchased from Sigma-Aldrich. The 3D model cell culture medium (DermaLife K Keratinocyte Medium) was purchased from Lifeline Cell Technology.

### N/TERT-2G keratinocytes cell culture

Frozen N/TERT-2G keratinocytes were initially thawed in a 37 °C water bath and resuspended in pre-warmed EpiLife/HKGS medium, then centrifuged at 300 g for 5 min at room temperature and resuspended in the EpiLife/HKGS medium. The cell suspension was then transferred into cell culture T75 flasks, with the medium changed every two days until 70-90% confluency was reached in each flask. Then, the cells were passaged by trypsinization with 5 mL of Trypsin-EDTA for 3 min at 37 °C, and the reaction was stopped by adding 10 mL of 10% FBS/DMEM. The cells were centrifuged at 300 g for 5 min at room temperature, then washed twice with 1× DPBS. Cell concentration was measured by mixing 10 μL of cell suspension with an equal volume of trypan blue (Bio-Rad) and counting with an automated cell counter (TC20™, Bio-Rad).

### CRISPR-Cas9 editing of N/TERT-2G keratinocytes

The crRNA (Alt-R^®^ CRISPR-Cas9 gRNA, IDT Technologies) sequences for targeting the start codon (ATG) of CYP11B1 were designed through CHOPCHOP (v3) ^[51]^; the crRNA sequences for a large deletion between exons were pre-designed and obtained from IDT Technologies (Table S1). The electroporation was conducted using the Neon System Kit (Neon™ Transfection System 100 μL Kit, Thermo Fisher Scientific) and Neon Transfection System (Thermo Fisher Scientific).

Initially, equal molar concentrations (40 μM) of each crRNA and tracrRNA (Alt-R® CRISPR-Cas9 tracrRNA, IDT Technologies) suspension were separately mixed and heated at 95 °C for 5 min to form the crRNA-tracrRNA duplex. Cas9 enzyme (Alt-R S.p. HiFi Cas9 Nuclease V3, IDT Technologies) suspension was prepared at a 30 μM concentration by dilution and mixed with the resuspension buffer T (Neon System Kit). Then, the ribonucleoprotein (RNP) complexes were prepared by mixing 180 pmol of crRNA-tracrRNA with 60 pmol of Cas9 and incubated at room temperature for 15 min. The RNP complexes for all sets of crRNAs and 180 pmol of enhancer (Alt-R Cas9 Electroporation Enhancer, IDT Technologies) were added to N/TERT-2G keratinocytes, which were resuspended in buffer R (Neon System Kit). Subsequently, the cell-RNP suspension was pipetted into the Neon tip (Neon System Kit) and inserted into the pipette station containing the electrolytic buffer (E2, Neon System Kit). The cells were electroporated at 1700 V with a 20 ms pulse width using the Neon Transfection System. After the electroporation, the cells were transferred into prewarmed EpiLife/HKGS medium and incubated in a 6-well plate for 48 h for further examination and monoclonal selection.

### DNA extraction, PCR and agarose gel electrophoresis

The CRISPR-Cas9-edited N/TERT-2G keratinocytes were cultured and expanded to a T25 flask, and 1 × 10⁵ to 1 × 10⁶ cells were then taken for genomic DNA extraction utilizing the QIAamp DNA Mini Kit (QIAGEN). The extracted DNA suspension was measured for quality and quantity using a NanoDrop One spectrophotometer (Thermo Fisher Scientific). Then, the edited region was amplified by PCR, and each reaction contained 25 μL of PCR Master Mix (K0171, Thermo Fisher Scientific), 5 μL of PCR primer mix (Table S2) at 10 μM, 200 ng DNA with a volume of 6 μL, and 14 μL of nuclease-free DI water. PCR amplification was done using a Thermo Cycler (Applied Biosystems) with the following setup: initial denaturation at 95 °C for 2 min, 38 cycles of denaturation-extension (95 °C for 30 s, 56 °C for 30 s, and 72 °C for 90 s), final extension at 72 °C for 10 min, and hold at 4 °C. Then, 20 μL of the amplified DNA suspension was loaded onto a 1.5% agarose gel containing GelRed Nucleic Acid Stain (Sigma-Aldrich) and run in 1× TAE buffer at 100 V, followed by imaging the gel on a ChemiDoc^TM^ MP imaging system (Bio-Rad).

### Monoclonal selection by Sanger sequencing and ICE analysis

The selected CRISPR-Cas9 KO keratinocyte pool was first resuspended and diluted to a final concentration of 0.8 cells per 100 μL of EpiLife/HKGS medium, then transferred into 96-well plates for single-cell culture. The wells containing a single cell clone were used for further expansion and monoclonal selection. After a series of expansions, keratinocytes were separated from the 12-well plate to ensure enough DNA could be extracted and amplified. Then, Sanger sequencing data was obtained by analyzing the extracted DNA in an ABI 3730 PRISM^®^ DNA Analyzer. The Sanger sequencing data of monoclonal groups were subsequently analyzed against the WT sequence using the ICE analysis tool (EditCo, Synthego). The sequences of both CYP11B1 KO and WT were aligned and analyzed in Benchling.

### Fabrication of the 3D epidermal model

As described in previous research ^[47]^, the *in-vitro* 3D epidermal model fabrication was initiated by coating the cell culture inserts with 2.5% collagen in 1× DPBS for 45 min at 4 °C. The WT or CYP11B1 KO monoclonal (B5 and DB4 clones) N/TERT-2G keratinocytes were resuspended in pre-warmed 3D model cell culture medium (DermaLife K Keratinocyte Medium) containing 1.5 mM CaCl₂. Then, the collagen solution was removed after coating, and 300 μL of cell suspension at a concentration of 1 × 10⁶ cells/mL was added to each cell culture insert. The cell culture inserts were placed in 6-well plates and cultured in 2.5 mL of pre-warmed 3D model cell culture medium containing 1.5 mM CaCl₂ in each well for 3 days. Afterward, the medium was removed from the cell culture inserts for air-lifting, and the surrounding medium was replaced with 3D model cell culture medium containing 1.5 mM CaCl₂ and 50 μg/mL Vitamin C. The 3D models were cultured for 12 days for keratinocyte differentiation, with the medium changed every 2 days until harvest.

### UVB irradiation

After 12 days of keratinocyte differentiation, the 3D models were cultured in a 3D model cell culture medium containing 1.5 mM CaCl₂ and 50 μg/mL Vitamin C, in the absence of cortisol for 4 h. Then, the 3D models were placed in a 24-well plate containing 1× DPBS and irradiated for 1250 s with two narrowband UVB lamps (PL-S 9W/01 G23, Philips) at an irradiance of 0.8 mW/cm², corresponding to a delivered dose of 1000 mJ/cm², as measured with a UV-AB radiometer (Extech Instruments). After UVB irradiation, the 3D models were transferred to new 6-well plates and cultured in 3D model cell culture medium containing 1.5 mM CaCl₂ and 50 μg/mL Vitamin C, in the absence of cortisol. The 3D models were cultured for an additional 24 h for sample collection and further analysis.

### Immunofluorescence analysis

The harvested 3D models were removed and cut from the cell culture inserts with disposable scalpels. The cut pieces of 3D models were fixed in 4% formaldehyde (VWR chemicals) at 4 °C overnight and transferred into embedding cassettes, followed by a graded ethanol dehydration and paraffin embedding in an Epredia Excelsior AS tissue processor. Subsequently, the paraffin-embedded 3D models were sectioned at a thickness of 7 μm using a microtome (Microm HM355S, Thermo Fisher Scientific) and mounted onto Superfrost™ Plus microscope slides (Epredia Netherlands B.V.).

The microscope slides were heated at 70 °C for 30 min and then immersed in Histolab-clear (Histolab) twice for 15 min each to remove the paraffin. This was followed by rehydration through a series of 100% isopropanol (twice), 96% ethanol, 70% ethanol, and deionized water for 5 min each. For antigen retrieval, the microscope slides were placed in a 2.1 g/L citric acid monohydrate solution (adjusted to pH 6.0 with 10 M NaOH) and heated in a microwave oven for 8 min. After cooling slowly under running tap water, the tissue sections were blocked for 1 hour at room temperature in an antibody dilution buffer (Abcam) containing 10% donkey serum (Sigma-Aldrich). The tissue sections were then incubated with primary antibodies (Table S3) overnight at 4 °C, washed 3 times with 1× PBS, and subsequently incubated with secondary antibodies for 1 h at room temperature, protected from light. In the end, the samples were mounted using ProLong Diamond Antifade Mountant (Thermo Fisher Scientific). Immunofluorescence imaging of the tissue sections was performed using a Zeiss LSM980-Airy confocal microscope with identical acquisition settings (laser power, gain and exposure) for all samples within an experiment, and the data were analyzed using Zeiss Zen Lite and ImageJ software. For quantification of CYP11B1 signal, the epidermal area of each section was outlined as a region of interest and the integrated density of the CYP11B1 channel was measured in ImageJ after background subtraction; each section from three independently generated models were analyzed per group.

### Scanning electron microscopy analysis

For the scanning electron microscopy (SEM) analysis, the 3D epidermal models were fixed overnight at 4 °C in a mixture containing 0.25% glutaraldehyde and 4% formaldehyde. Subsequently, the samples were rinsed with ultrapure water (MilliQ water) and dehydrated through a graded series of ethanol solutions. Critical point drying was performed using an EM CPD300 dryer (Leica), and the samples were then mounted onto specimen stubs using conductive carbon adhesive. A 10 nm-thick platinum layer was sputter-coated onto the sample surfaces using a Q150T ES sputter coater (Quorum) system. Imaging was conducted on a Gemini Ultra 55 field-emission scanning electron microscope (Zeiss) at an accelerating voltage of 3 kV, utilizing an SE2 detector. Image analysis was performed using ImageJ software.

### Histological analysis

Following the previously described protocol for the preparation of tissue sections for immunofluorescence analysis, the microscope slides were heated at 70 °C for 30 min to remove the paraffin. Subsequently, the microscope slides were immersed in Histolab-clear twice (15 min each time), followed by sequential immersion in 100% ethanol (twice), 96% ethanol, 70% ethanol, and deionized water, with each step lasting 4 min. The rehydrated tissue sections were stained with Mayer’s Hematoxylin (HTX) at room temperature for 10 min, rinsed in running tap water for 10 min, and then stained with 0.2% eosin for 2 seconds. After a thorough rinse under running tap water, the stained tissue sections were dehydrated (by sequential immersion in a graded series of ethanol concentrations, followed by Histolab-clear) and finally mounted using EcoMount. Histological images were acquired using a bright-field microscope (Nikon Eclipse Ni) and processed with ImageJ software.

### RNA isolation and RT-qPCR analysis

Messenger RNA (mRNA) was isolated from the 3D epidermal models to analyze the transcription and expression of target genes. First, the 3D epidermal models were washed twice with 1× DPBS, cut from the cell culture inserts as previously described, and transferred into RNase-free Eppendorf tubes. Subsequently, 500 μL of TRIzol reagent (Invitrogen) was added to each tube, followed by incubation at room temperature for 5 min; then, 100 μL of chloroform was added, and the samples were incubated for an additional 2 min. The tissue suspensions were centrifuged at 12,000 rcf for 15 min at 4 °C. Following centrifugation, the upper phase solution was transferred into a new Eppendorf tube containing 1.5 μL of GlycoBlue (Invitrogen), and 250 μL of isopropanol was added. After a 10-minute incubation, the samples were centrifuged at 12,000 rcf for 10 min at 4 °C to precipitate the RNA. The RNA pellet was then washed twice with 75% ethanol, air-dried, and resuspended in deionized water. The RNA resuspension was measured for quality and quantity using a NanoDrop One spectrophotometer (Thermo Fisher Scientific).

The cDNA synthesis was performed using the High-Capacity cDNA Reverse Transcription Kit (Applied Biosystems) with an input of 500 ng of mRNA per sample, followed by RT-PCR amplification. The amplification protocol consisted of incubation at 25 °C for 10 min, 37 °C for 120 min, 85 °C for 5 min, and hold at 4 °C. Subsequently, 10 ng of cDNA (in a volume of 4.5 μL) was mixed with 0.5 μL of a qPCR primer mix (Table S4 and S5), and 5 μL of either TaqMan™ Gene Expression Master Mix (TaqMan) or SYBR™ Green Universal Master Mix (Applied Biosystems). Real-time quantitative PCR (qPCR) assays were performed using the QuantStudio 6/7 (Thermo Fisher Scientific). Relative mRNA levels were calculated using the 2^−ΔΔCt^ method with GAPDH as the reference gene and the WT control group as the calibrator. Data plotting and analysis were conducted using OriginLab Software.

### Statistical analysis

Data are presented as mean ± standard deviation of (at least three) independently generated 3D epidermal models per group (biological replicates); technical replicates were averaged before analysis. Immunofluorescence integrated density was compared between groups from three different 3D epidermal models, and RT-qPCR 2^−ΔΔCt^ values were compared across five different 3D epidermal models within each group. All statistical analyses and data visualization were performed in OriginLab Software. Quantification comparison data plots were generated using the Paired Comparison Plot App to visualize the distribution and variability of the experimental data. Pairwise comparisons between experimental groups were performed using *Tukey’s* multiple comparison test, as implemented in the App. Differences were considered statistically significant at *P* ≤ 0.05 (\**P* ≤ 0.05, \*\**P* ≤ 0.01, \*\*\**P* ≤ 0.001).

## Acknowledgements

The N/TERT-2G human keratinocyte cell line was obtained from Jos P.H. Smits and Ellen H. van den Bogaard (Department of Dermatology, Radboud Research Institute for Medical Innovation, Radboud University, Nijmegen, The Netherlands) on behalf of James Rheinwald (Department of Dermatology, Brigham and Women’s Hospital, Harvard Medical School, Boston, MA, USA). We thank Histological Core Facility, the Biomedicum Imaging Core Facility (BIC), and the Electron Microscopy Core Facility (EMil) at Karolinska Institutet, for sample preparation and measurements. We thank KI Gene at Karolinska Institutet for support with the QuantStudio 6/7 real-time qPCR equipment and Sanger DNA sequencing. O.P. acknowledges Swedish Research Council (Vetenskapsrådet, 2022-01436) for financial support.

## Author Contributions

S.X.: conceptualization, methodology, investigation, formal analysis, visualization, writing – original draft. K.D.M.: methodology, investigation, writing – review and editing. G.N.: methodology, writing – review and editing. O.P.: conceptualization, supervision, funding acquisition, project administration, writing – review and editing. All authors have read and approved the final version of the manuscript.

## Data Availability Statement

The data that support the findings of this study are available from the corresponding author upon request. The Sanger sequencing traces and ICE analysis outputs for the wild-type and edited clones, together with the uncropped gel images, are provided as Supporting Information.

## Supporting Information

Supporting Information is available from the author.

Received: ((will be filled in by the editorial staff))

Revised: ((will be filled in by the editorial staff))

Published online: ((will be filled in by the editorial staff))

