## Supporting Information for "Genetic disruption of cutaneous de novo glucocorticoid synthesis: a CRISPR-Cas9 CYP11B1-knockout human epidermal model"

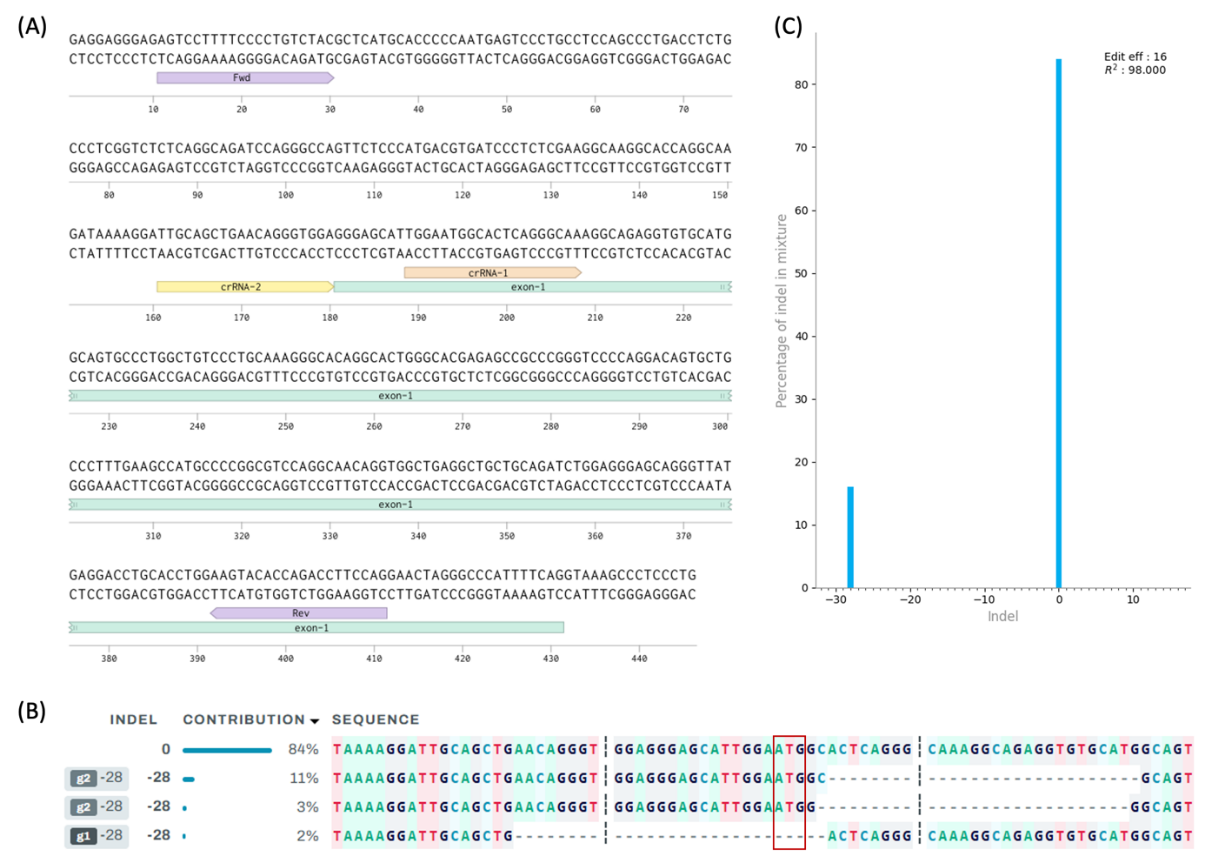

**Figure S1**. *Strategy I:* indels targeting the start codon (ATG) of CYP11B1. (A) Schematic of crRNA and primers design. (B) Sanger sequencing data of ATG KO pool aligned with WT, analyzed by ICE. (C) Indels and their relative prevalence for the ATG KO pool, calculated by ICE.

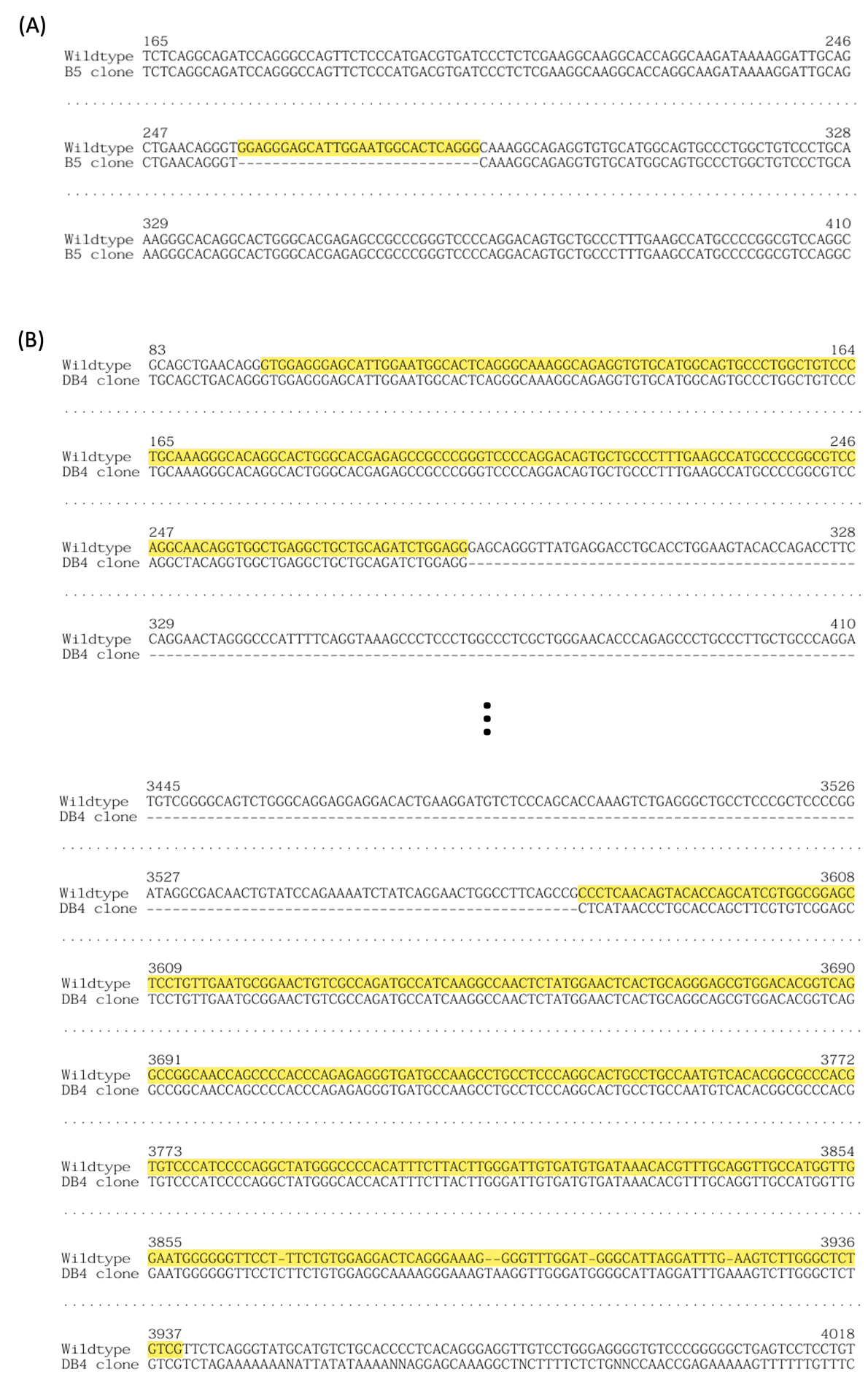

**Figure S2**. Genomic DNA sequence aligned with WT using Benchling. (A) B5 monoclonal group sequence, the indel mutation region is highlighted. (B) DB4 monoclonal group sequence; the retained region is highlighted.

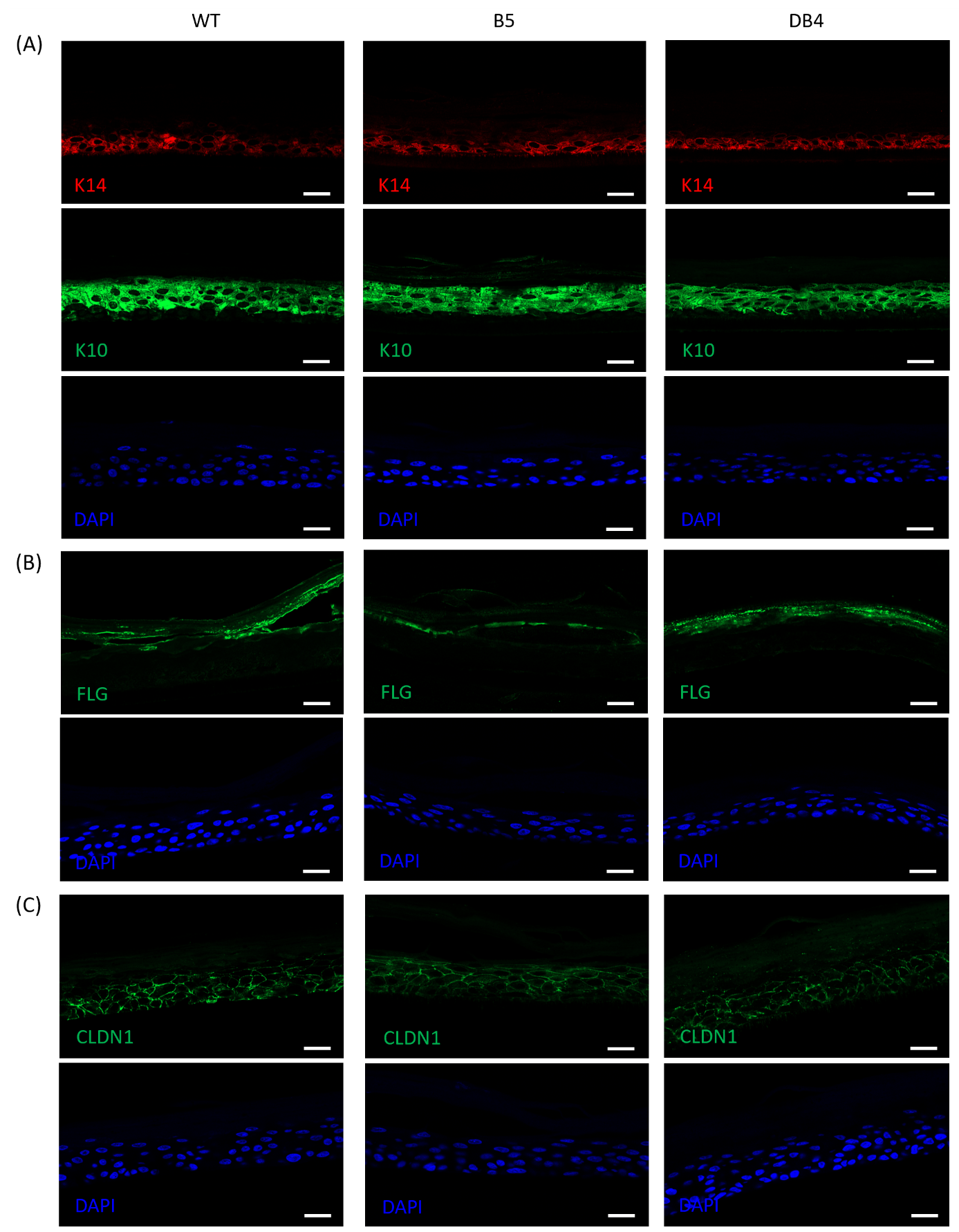

**Figure S3**. Split channels for each immunofluorescent stained 3D epidermal sections. (A) Key structural biomarkers. Red: K14, Green: K10, Blue: DAPI. (B) Key differentiation biomarker. Green: FLG, Blue: DAPI. (C) Key intercellular biomarker. Green: CLDN1, Blue: DAPI. Scale bars: 20 μm.

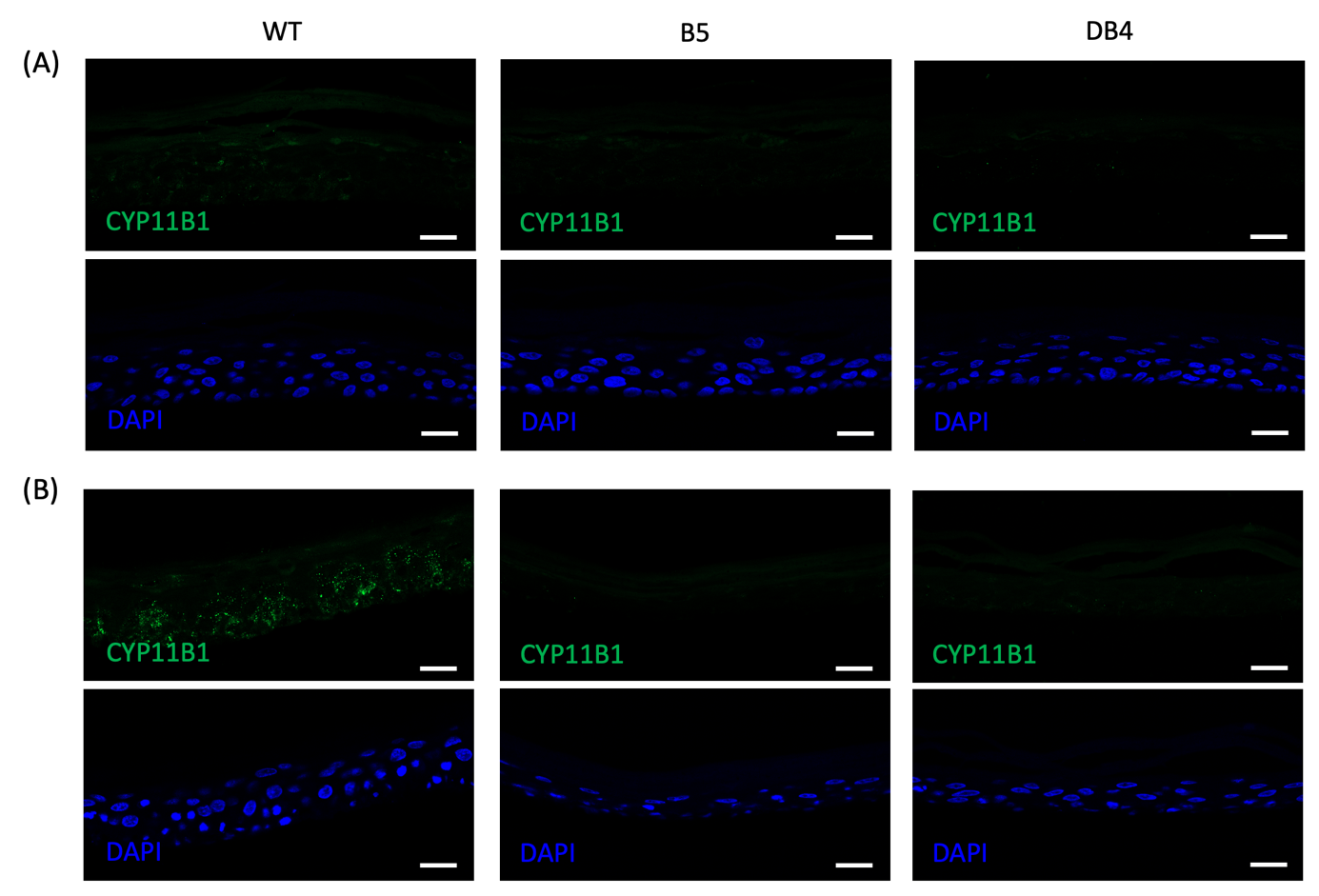

**Figure S4.** Split channels for CYP11B1 immunofluorescent stained 3D epidermal sections. (A) Control group, 3D epidermal models without any treatment. (B) UVB-treated 3D epidermal models. Green: CYP11B1, Blue: DAPI. Scale bars: 20 μm.

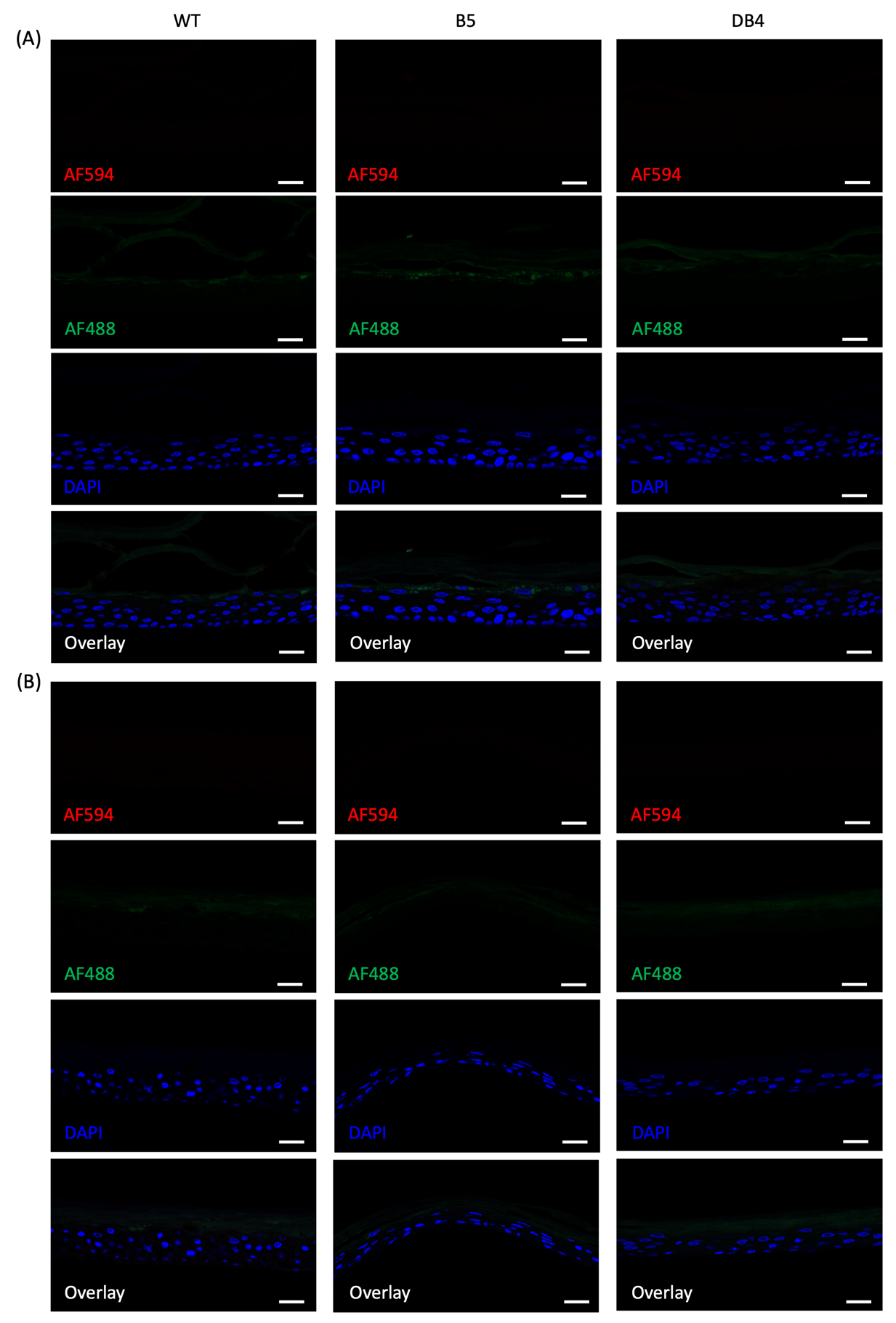

**Figure S5.** Split channels for negative immunofluorescent stained 3D epidermal sections, primary antibody omitted and replaced with antibody diluent buffer. (A) Control group, 3D epidermal models without any treatment. (B) UVB-treated 3D epidermal models. Blue: DAPI. Scale bars: 20 μm.

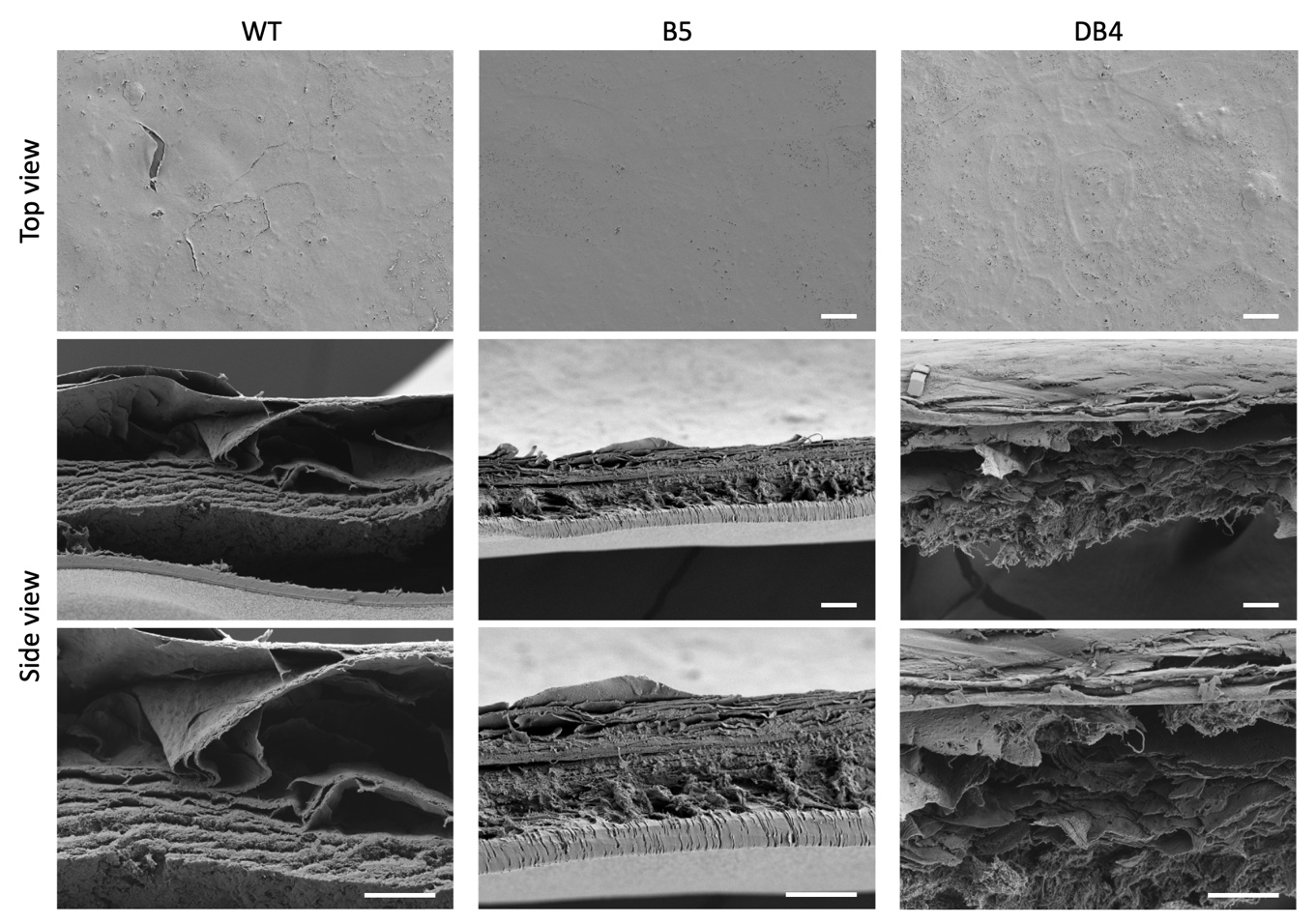

**Figure S6.** Scanning electron microscopy images of the 3D epidermal models after UVB treatment. Scale bars: 20 μm.

**Table S1.** **crRNA sequence design**.

| Strategy | crRNA (IDT id.) | crRNA sequence (5’ – 3’) | PAM Site |
| --- | --- | --- | --- |
| I. Indels on ATG | crRNA-1 | Forward: TGGAATGGCACTCAGGGCAA | AGG |
|  | crRNA-2 | Forward: TTGCAGCTGAACAGGGTGGA | GGG |
| II. Large deletion | crRNA-1 (Hs.Cas9.CYP11B1.1.AE) | Forward: GGGTTATGAGGACCTGCACC | TGG |
|  | crRNA-2 (Hs.Cas9.CYP11B1.1.AD) | Reverse: GAGGAACCTCTGCACAGCGT | TGG |
|  | crRNA-3 (Hs.Cas9.CYP11B1.1.AA) | Reverse: ACGATGCTGGTGTACTGTTG | AGG |

**Table S2. CYP11B1 PCR primer sequence for Sanger sequencing and monoclonal selection.**

| Strategy | Forward Primer (5’ – 3’) | Reverse Primer (5’ – 3’) |
| --- | --- | --- |
| I. Indels on ATG | AGTCCTTTTCCCCTGTCTAC | CTGGAAGGTCTGGTGTACTT |
| II. Large deletion (1^st^ screening) | ACAGCCCCCACTCCATGGCA | GTCTGGCACCACCTCACCC |
| II. Large deletion (2^nd^ screening) | TGATCCCTCTCGAAGGCAAG | ACATGCATACCCTGAGAACG |

**Table S3. The primary antibodies and secondary antibodies utilized for the immunofluorescent stain.**

The anti-CYP11B1 Antibody, clone 80-7 was obtained from Sigma-Aldrich. All the other antibodies were obtained from Abcam. The antibodies were all diluted with antibody diluent buffer (ab64211, Abcam).

| Antibodies | Cat. no. | Target | Host | Dilution |
| --- | --- | --- | --- | --- |
| Anti-Cytokeratin 14 antibody | ab7800 | K14 | Mouse | 1:100 |
| Anti-Cytokeratin 10 antibody | ab76318 | K10 | Rabbit | 1:100 |
| Anti-Filaggrin antibody | ab81468 | Filaggrin | Rabbit | 1:100 |
| Anti-Claudin 1 antibody | ab211737 | Claudin 1 | Rabbit | 1:100 |
| Anti-CYP11B1 Antibody, clone 80-7 | MABS502 | CYP11B1 | Rat | 1:50 |
| Donkey Anti-Mouse IgG H&L (Alexa Fluor® 594) | ab150108 | Mouse IgG |  | 1:1000 |
| Donkey Anti-Rabbit IgG H&L (Alexa Fluor® 488) | ab150073 | Rabbit IgG |  | 1:1000 |
| Donkey Anti-Rat IgG H&L (Alexa Fluor® 488) | ab150153 | Rat IgG |  | 1:200 |

**Table S4. Quantitative PCR Primer Sequences, TaqMan-based detection.**

| Gene | IDT id. | Primer sequence (5’ – 3’) | Location |
| --- | --- | --- | --- |
| CYP11B1 | Hs.PT.58.40046582.g | Forward: AACCTTCTTGCTGTGTCCTC  Reverse: TGCCCTCTTGAATGGAACTG | exon 9 |
| GAPDH | Hs.PT.39a.22214836 | Forward: ACATCGCTCAGACACCATG  Reverse: TGTAGTTGAGGTCAATGAAGGG | exon 2-3 |

**Table S5. Quantitative PCR Primer Sequences, SYBR Green-based detection.**

| Genes | Full name | Primer sequence (5’ – 3’) |
| --- | --- | --- |
| HSD11B1 | Hydroxysteroid 11-beta dehydrogenase 1 | Forward: CTGCCTGCTTAGGAGGTTGTA  Reverse: GTAGTAGGCCATGAAGAGCCC |
| HSD11B2 | Hydroxysteroid 11-beta dehydrogenase 2 | Forward: GTGCTCATCACCGGCTGT  Reverse: GCTGTTCAACTCCAATACGG |
| GAPDH | Glyceraldehyde-3-phosphate dehydrogenase | Forward: GGTGTGAACCATGAGAAGTATGA  Reverse: GAGTCCTTCCACGATACCAAAG |
